# CRISPR-DT: designing gRNAs for the CRISPR-Cpf1 system with improved target efficiency and specificity

**DOI:** 10.1101/269910

**Authors:** Houxiang Zhu, Chun Liang

## Abstract

The CRISPR-Cpf1 system has been successfully applied in genome editing. However, target efficiency of the CRISPR-Cpf1 system varies among different gRNA sequences. We reanalyzed the published CRISPR-Cpf1 gRNAs data and found many sequence and structural features related to their target efficiency. Using machine learning technology, a SVM model was created to predict target efficiency for any given gRNAs. We have developed the first web service application, CRISPR-DT (CRISPR DNA Targeting), to help users design optimal gRNAs for the CRISPR-Cpf1 system by considering both target efficiency and specificity. CRISPR-DT is available at http://bioinfolab.miamioh.edu/CRISPR-DT.

## Background

The CRISPR-Cas9 (Clustered Regularly Interspaced Short Palindromic Repeats - CRISPR associated protein 9) system is a prokaryotic immune system, which has been widely applied in eukaryotic genome editing. The basic mechanism of CRISPR-Cas9 editing is that a single guide RNA (gRNA) guides Cas9 to a DNA target sequence, then Cas9 cleaves double strands at the same position. The DNA double-strand breaks can be repaired either by non-homologous end joining (NHEJ), leading to a gene knockout, or by homology directed repair (HDR), generating a gene knockin. In 2013, the CRISPR-Cas9 system was first demonstrated as a tool for eukaryotic genome editing [1, 2]. Afterwards, many studies have successfully applied CRISPR-Cas9 to engineer genomes of different organisms [3–11]. In addition, researchers found that CRISPR-Cas9 can even be used to correct genetic diseases [12, 13] and develop cancer models [6, 14–16]. But the CRISPR-Cas9 system still has some shortcomings, such as high off-target effect [17, 18]. The CRISPR-Cpf1 system was first characterized in 2015, and it is a class 2 CRISPR-Cas system with a single Cas protein to mediate cleavage like CRISPR-Cas9 [19]. Researchers found that CRISPR-Cpf1 has some characteristics distinct from those of CRISPR-Cas9 [19]. First, Cpf1 has RNase III activity for processing the precursor CRISPR RNA (pre-crRNA) into mature crRNAs [20, 21]. Second, Cpf1 needs only one crRNA for cleavage, whereas Cas9 requires both the crRNA and trans-activating crRNA (tracrRNA) [19]. Third, Cpf1 recognizes a T-rich protospacer adjacent motif (PAM), which is at the 5’ end of the protospacer sequence [19]. Fourth, Cpf1 cleaves double-strand DNA via a staggered cut that generates two sticky ends, and the cleavage sites are further away from the PAM [19]. Recently, the CRISPR-Cpf1 system has been successfully applied for genome editing in various organisms. Studies showed that CRISPR-Cpf1 is highly specific in mammals [22–25]. Compared with Cas9, Cpf1 induced less off targets in human cells [23]. Cpf1 also has high target specificity in plant cells [26–29]. In addition, CRISPR-Cpf1 has been used to genetically engineer some important bacteria, such as *Corynebacterium glutamicum* [30] and *Cyanobacteria* [31], whose genomes cannot be edited by the CRISPR-Cas9 system probably due to Cas9 toxicity in these bacteria [30, 32]. Moreover, researchers demonstrated that CRISPR-Cpf1 is highly suitable for multiplex gene editing [21, 33]. CRISPR-Cas9 can also be used in multiplex gene editing, but it requires larger constructs or delivery of several plasmids [34–37], which might cause problems in multiplex gene editing [21]. Additionally, the CRISPR-Cpf1 system has been applied in correction of genetic mutations [38, 39], which shows great therapeutic potential. Furthermore, the inactive Cpf1 can be used for gene repression [26, 40]. All these studies demonstrate that the CRISPR-Cpf1 system has many advantages over CRISPR-Cas9, and it is a powerful tool for genome editing.

Target efficiency and specificity are the two most important aspects for genome editing. However, target efficiency of the CRISPR-Cpf1 system varies among different gRNA sequences [41], just like the CRISPR-Cas9 system [7, 8, 42–46]. Kim and his colleagues attempted to find out gRNA features that are related to target efficiency of CRISPR-Cpf1 [41]. They measured activities of 1251 gRNA sequences for the CRISPR-Cpf1 system and developed an algorithm to predict target efficiency [41]. In our study, we reanalyzed this public data by comparing the most efficient gRNAs (top 10% in activity ranking) with the least efficient gRNAs (bottom 10%). The most distinct gRNA features related to target efficiency can be identified by excluding gRNAs with modest efficiency, which has been demonstrated as a great strategy in previous studies [47, 48]. We found many novel gRNA sequence and structural features that are highly beneficial to target efficiency of CRISPR-Cpf1. Using machine learning technology, we created a support vector machine (SVM) model to predict target efficiency by combining all the important features selected by Random Forest. Previous research showed that the SVM had robust performance in predicting target efficiency for the CRISPR-Cas9 system [48]. Random Forest can rapidly screen optimal features [49], which has been applied in various biological studies [50–52]. We have developed the first web service application, CRISPR-DT, to help users design gRNAs for the CRISPR-Cpf1 system by considering both target efficiency and specificity.

## Results

### Sequence features of efficient and inefficient gRNAs

#### Position-specific nucleotide composition

Fig. 1 shows that within the 23-nt gRNA sequence, uracil (U) is extremely disfavored at the first position of efficient gRNAs, which is immediately adjacent to the PAM sequence (*P* = 2.44E-14). But guanine(G) and cytosine (C) are strongly favored at the first position (*P* = 6.96E-06 and *P* = 1.14E-04). In the last position, efficient gRNAs prefer cytosine (*P* = 5.42E-06), but not guanine (*P* = 3.16E-04). Guanine is especially disfavored at the end of efficient gRNAs (position 18 - 23). Overall, within the 23-nt RNA sequence, efficient gRNAs favor adenine (A) but disfavor guanine except at the first position, probably due to the first position is adjacent to the PAM sequence. Position-specific dinucleotides were also analyzed by comparing efficient and inefficient gRNAs (Additional file 1: Table S1). For the first two positions of efficient gRNAs, CC and GG are favored with enrichment ratios of 4.57 (*P* = 1.12E-05) and 3.00 (*P* = 4.94E-03), whereas UC, UU, and UG are disfavored with enrichment ratios of 0.05 (*P* = 1.51E-05), 0.00 (*P* = 7.08E-05), and 0.08 (*P* = 9.91E-04), respectively.

**Fig. 1.**
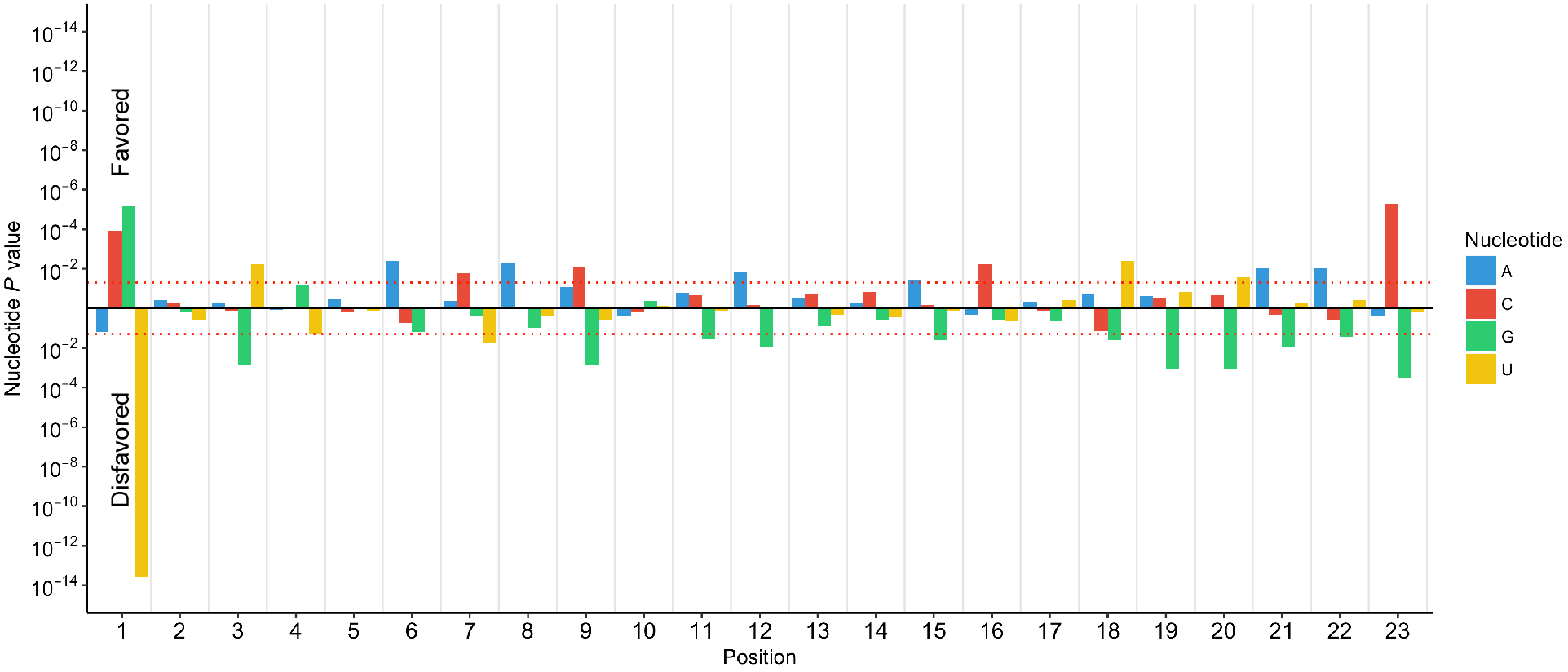
*P*-values of single nucleotides at each position of the 23-nt gRNA sequence. The Y axis indicates whether a single nucleotide is favored or disfavored by efficient gRNAs. The red lines are used as the cutoff for significance (*P* = 0.05).

#### Position-nonspecific nucleotide composition

Table 1 shows that the average counts of guanine for efficient and inefficient gRNAs are 5.66 and 7.53, respectively, within the 23-nt gRNA sequence (*P* = 2.59E-08), which indicates efficient gRNAs disfavor guanine. But compared with inefficient gRNAs, efficient gRNAs have a preference for adenine (5.50 versus 4.30, *P* = 1.57E-06) and cytosine (7.04 versus 6.01, *P* = 1.24E-03). The most significant position-nonspecific dinucleotide count is the GG count (1.30E-06), which greatly decreased in efficient gRNAs with an enrichment ratio of 0.59. This finding is consistent with previous study of CRISPR-Cas9 [48]. The position-nonspecific trinucleotide and tetranucleotide counts were also calculated (Additional file 1: Table S2). UGG is the most significant trinucleotide for efficient and inefficient gRNAs with average counts of 0.32 and 0.75, respectively (*P* = 1.83E-05). The average count of GGG is significantly decreased in efficient gRNAs with an enrichment ratio of 0.57 (*P* = 8.09E-03), which is consistent with previous CRISPR-Cas9 study [48]. CCCA is the most significant tetranucleotide for efficient and inefficient gRNAs with average counts of 0.17 and 0.02, respectively (*P* = 3.84E-04). Previous work has demonstrated that GGGG within the 23-nt gRNA sequence can cause poor CRISPR-Cas9 activity because GGGG badly affects DNA oligo synthesis, and can form a secondary structure called guanine tetrad in the gRNA sequence, which makes gRNAs difficult to bind to target sequences [48]. Consistently, for the CRISPR-Cpf1 system, we found there are more GGGG in inefficient gRNAs than that in efficient gRNAs with an enrichment ratio of 0.21 (*P* = 4.48E-04). Research also showed that repetitive nucleobases (at least four A, four C, four G, or four U) are related to poor CRISPR-Cas9 activity [48]. Similarly, in our study we found more inefficient gRNAs contains repetitive nucleobases than efficient ones (0.30 versus 0.17, *P* = 1.65E-02). In addition, we examined UUU in the gRNA seed region, which are the 6 nucleotides in the 5’ PAM-proximal region [41], and found that more inefficient gRNAs contains UUU in the seed region than efficient ones with an enrichment ratio of 0.29 (*P* = 9.07E-02), which is consistent with previous CRISPR-Cas9 research [48].

**Table 1.** Significant position-nonspecific mononucleotide and dinucleotide counts within the 23-nt gRNA sequences

| Mono- and Dinucleotides | Efficient gRNAs | Inefficient gRNAs | Enrichment Ratio<sup>a</sup> | *P*-value<sup>b</sup> |
| --- | --- | --- | --- | --- |
| Count_G | 5.66 | 7.53 | 0.75 | 2.59E-08 |
| Count_A | 5.50 | 4.30 | 1.28 | 1.57E-06 |
| Count_C | 7.04 | 6.01 | 1.17 | 1.24E-03 |
| Count_GG | 1.19 | 2.03 | 0.59 | 1.30E-06 |
| Count_CG | 1.10 | 1.74 | 0.64 | 1.72E-05 |
| Count_AC | 2.14 | 1.48 | 1.44 | 9.63E-05 |
| Count_UA | 1.01 | 0.64 | 1.58 | 1.08E-03 |
| Count_AU | 0.79 | 0.49 | 1.62 | 1.23E-03 |
| Count_CU | 1.67 | 1.23 | 1.36 | 1.96E-03 |
| Count_CC | 1.68 | 1.22 | 1.37 | 3.88E-03 |
| Count_GC | 0.87 | 1.28 | 0.68 | 5.00E-03 |
| Count_AA | 0.91 | 0.60 | 1.52 | 5.49E-03 |
| Count_GU | 1.50 | 1.92 | 0.78 | 8.72E-03 |
| Count_UG | 1.30 | 1.66 | 0.78 | 1.28E-02 |
| Count_CA | 1.70 | 1.35 | 1.25 | 1.73E-02 |

<sup>a</sup>The enrichment ratio was calculated through dividing the average nucleotide counts of efficient gRNAs by that of inefficient gRNAs.
<sup>b</sup>The *P*-value was determined by Welch's t-test ( $P < 0.05$ ).

#### GC content

Researchers have demonstrated that the 23-nt gRNA sequence of CRISPR-Cpf1 should be divided into three regions: seed (6 nucleotides in the 5’ PAM-proximal region), trunk (12 nucleotides in the middle region), and promiscuous (5 nucleotides in the 3’ PAM-distal region) [41], which is illustrated in Additional file 1: Figure S1. The tolerant ability of mismatches gradually increases from the seed region to the promiscuous region. Here we separately compared GC content of the entire 23-nt gRNA sequence, seed region, trunk region, and promiscuous region between efficient and inefficient gRNAs. For the entire 23-nt gRNA sequence, inefficient gRNAs have higher GC content than efficient ones (0.59 versus 0.55, *P* = 3.54E-03). In addition, the GC content of inefficient gRNAs is higher than that of efficient ones in both trunk and especially promiscuous regions (0.58 versus 0.54, *P* = 7.41E-03 and 0.64 versus 0.52, *P* = 8.16E-06), but there is no big difference of GC content in the seed region. Since gRNAs with balanced GC content have higher target efficiency for the CRISPR-Cas9 system [8, 42, 53], in our study we consider GC content of 0.30 - 0.70 as normal, but greater than 0.70 or less than 0.30 as abnormal. Results showed that more efficient gRNAs contain normal GC content than inefficient ones (0.98 versus 0.89, *P* = 1.95E-03).

### Structural features of efficient and inefficient gRNAs

#### Minimum free energy

The secondary structure stability of the 23-nt gRNA sequence was determined by minimum free energy (MFE). Compared with efficient gRNAs, more inefficient gRNAs have lower MFE but fewer ones have higher MFE (Fig. 2). In addition, the average MFE of inefficient gRNAs is significantly lower than that of efficient ones (−3.89 versus −1.81, *P* = 1.10E-11), which means the secondary structure of inefficient gRNAs is more stable than that of efficient ones. Our finding indicates that nucleotide accessibility of the 23-nt gRNA sequence is strongly related to target efficiency, which is consistent with previous CRISPR-Cas9 study [48].

**Fig. 2.**
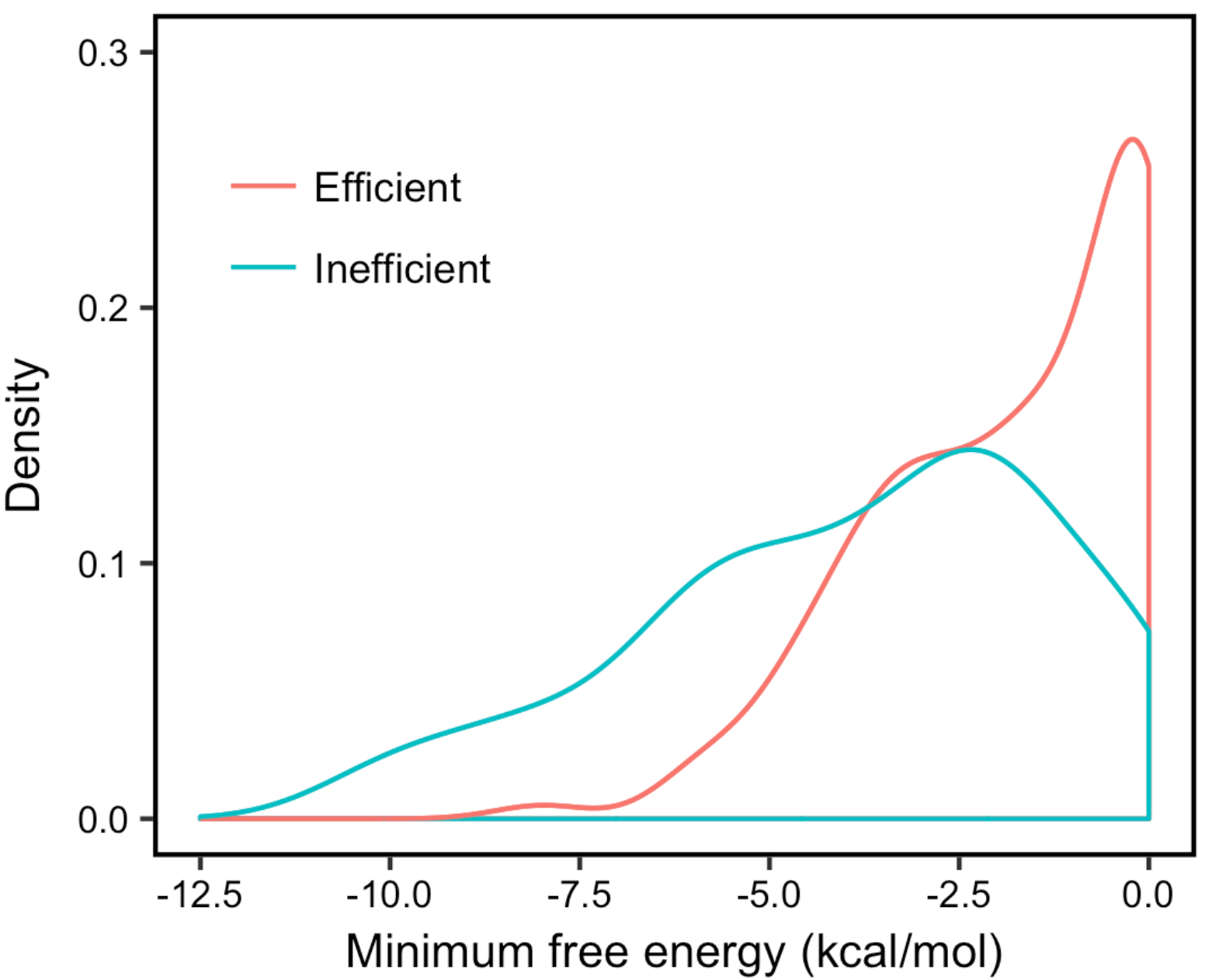
The distribution of efficient and inefficient gRNAs for minimum free energy.

#### Melting temperature

The melting temperature was calculated based on the DNA version of the gRNA sequence [54]. We separately compared melting temperatures of the entire 23-nt gRNA sequence, seed region, trunk region, and promiscuous region between efficient and inefficient gRNAs, which is a similar strategy as previous research [54]. For the entire 23-nt gRNA sequence, inefficient RNAs have a significantly higher average melting temperature than efficient ones (63.85 versus 61.34, *P* = 1.29E-06). Additionally, the average melting temperatures of inefficient gRNAs are higher than those of efficient ones in both trunk (38.03 versus 35.96, *P* = 5.37E-03) and promiscuous (−24.14 versus −31.84, *P* = 1.58E-07) regions, but not in the seed region.

### A SVM model to predict target efficiency

The top 10% and bottom 10% (250 total) gRNAs in activity ranking of the 1251 gRNAs were used to train and test our support vector machine (SVM) models. Since the full set of sequence and structural features was overdetermined, feature selection was performed before training SVM models. We used ten-fold cross validation to evaluate the SVM models and the receiver operating characteristic (ROC) curves are shown in Fig. 3. The average area under the curve (AUC) is 0.87, which means the SVM models have high performance in distinguishing efficient and inefficient gRNAs. We built a final SVM model by using all 250 gRNAs with the most important features. To further validate the SVM model,we compared our model with the previously published model [41], which was based on 104 independent gRNAs data (Fig. 4a). Our model showed to have a better performance than Kim et al.’s model (AUC = 0.76 versus 0.67) in predicting target efficiency using 104 independent gRNAs. Since research has demonstrated that TTTA is the preferred PAM sequence for the CRISPR-Cpf1 system relative to other PAM sequences [41], it’s very important to evaluate our model by only using gRNAs with the TTTA PAM. Fig. 4b shows our model has a better performance than Kim et al.’s model (AUC = 0.79 versus 0.44) in predicting target efficiency using 35 independent gRNAs with the TTTA PAM. Additionally, we evaluated our model based on gRNAs with TTTC and TTTG PAMs, separately (Additional file 1: Figure S2). Our model performs better at gRNAs with the TTTC PAM than Kim et al.’s model (AUC = 0.78 versus 0.71) and both models show similar results on gRNAs with the TTTG PAM (AUC = 0.71 versus 0.72). In summary, our SVM model has robust performance at predicting target efficiency.

**Fig. 3.**
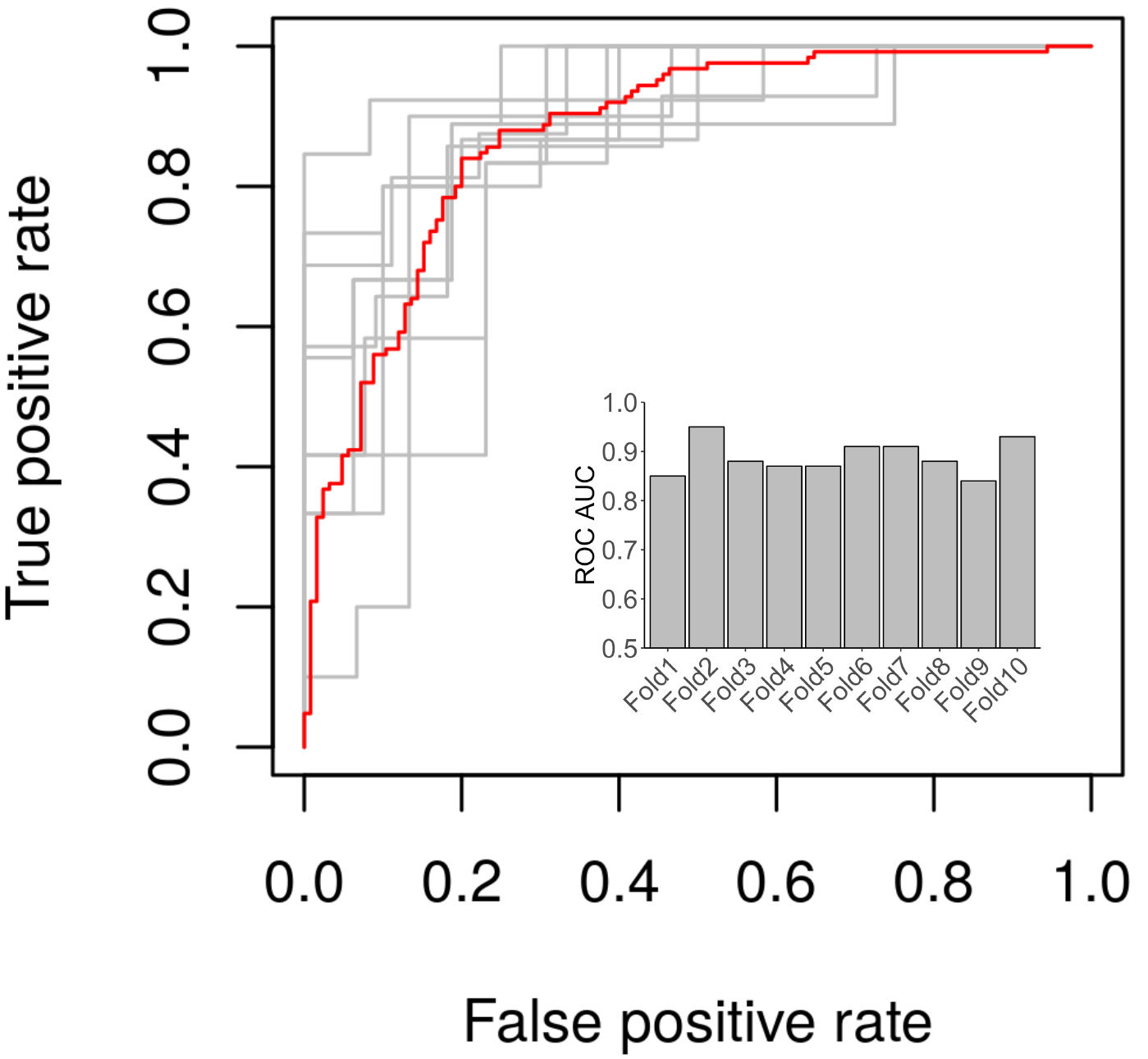
Performance evaluation of SVM models. Ten-fold cross validation was used to evaluate the SVM models. Each gray line indicates the ROC curve for each fold. The red line is the average ROCcurve. The bar graph indicates the AUC for each fold.

**Fig. 4.**
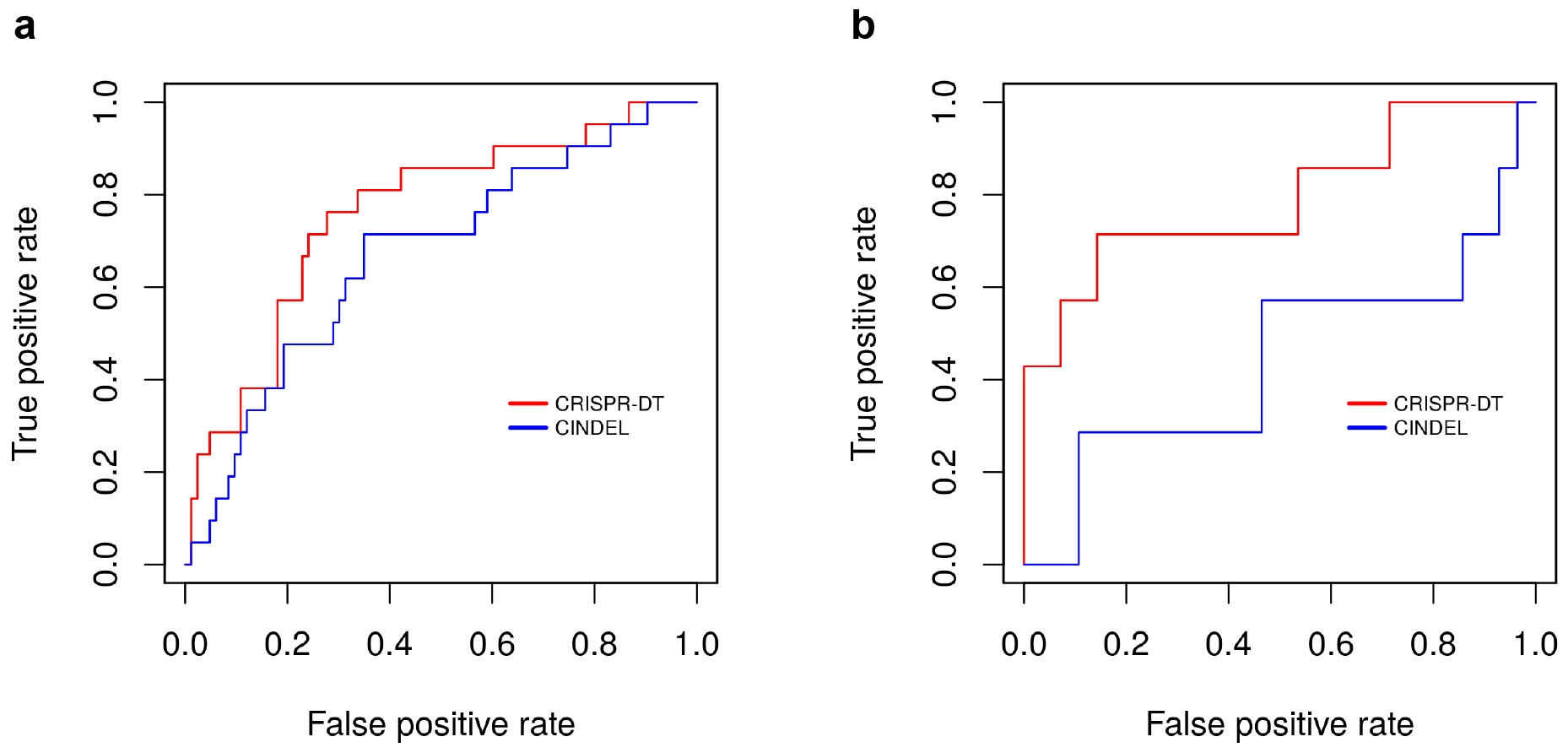
Validation of our final SVM model using independent experimental data. **a** ROC curves comparing the performance of our model (CRISPR-DT) and Kim et al.’s model (CINDEL)in predicting target efficiency using 104 independent gRNAs. **b** ROC curves comparing the performance of our model (CRISPR-DT) and Kim et al.’s model (CINDEL) in predicting target efficiency using independent gRNAs with the TTTA PAM.

### A web service application for improving target efficiency and specificity

Though the CRISPR-Cpf1 system is highly specific in human and plant cells, we still can use bioinformatics methods to improve target specificity at the greatest extent. By considering both target efficiency and specificity, we have developed a web service application, CRISPR-DT, to help users design optimal gRNAs for the CRISPR-Cpf1 system. In the setting page (Additional file 1: Figure S3), first, users input a DNA sequence they want to target in FASTA format. Second, users select a reference genome. Third, users set on-and off-target PAMs, respectively. Fourth, users choose an off-target setting. Then they can click “Find targets!” to run the program. In the result pages, Fig. 5a shows all the target candidates and relevant information, including the number of target sites within the entire genome, the number of exon targets within the entire genome, and the efficiency score of each target candidate. Users can rank target candidates by clicking column headers and choose optimal ones by considering both off-target effect and target efficiency. The number of target sites and the number of exon targets can be clicked to show the detailed information for each target site (Fig. 5b), including alignment information and the JBrowse [55] link, which can be clicked to display the target site in the background of genomic and transcript features (Fig. 5c).

**Fig. 5.**
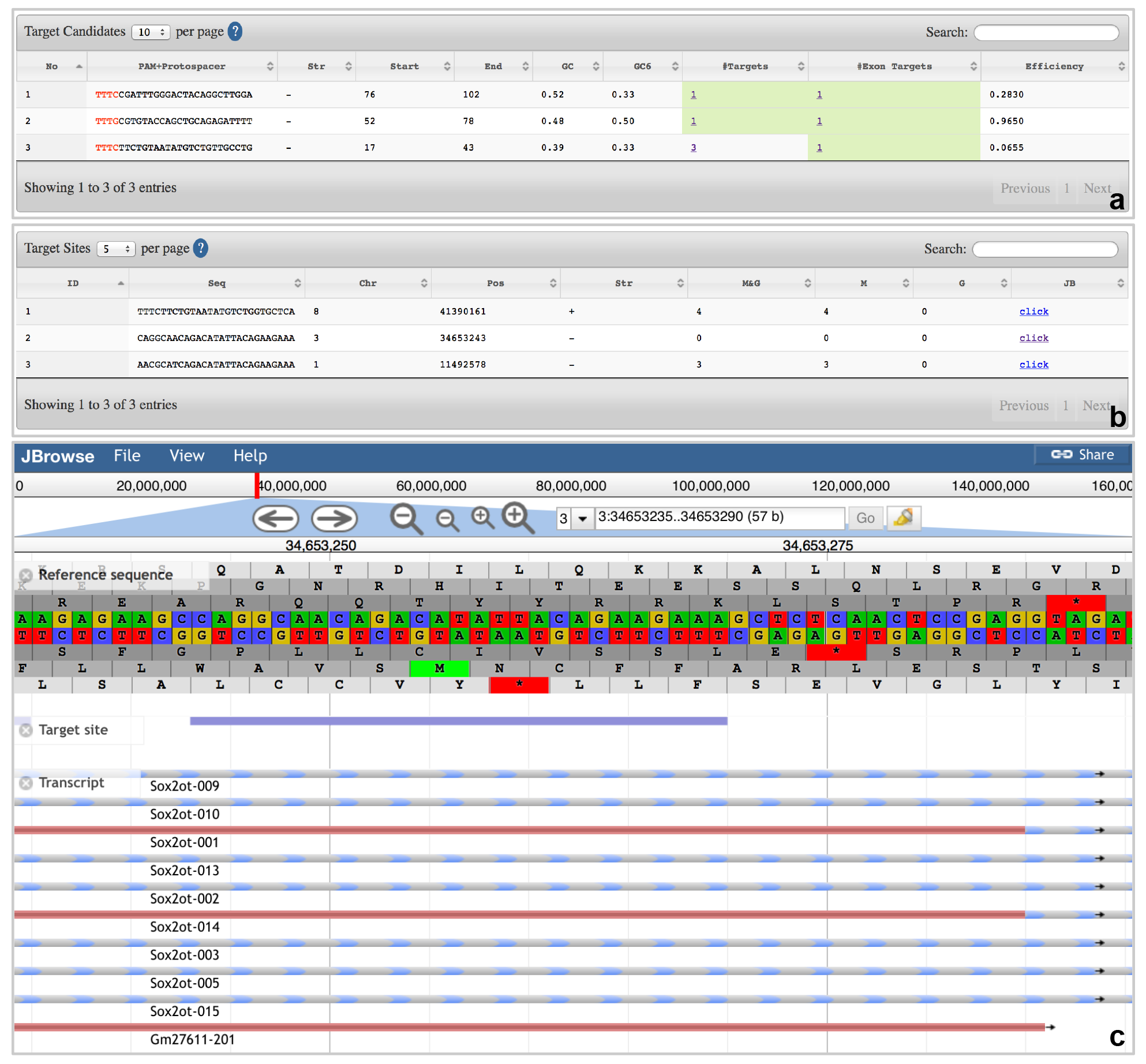
The result web interfaces of CRISPR-DT. **a** The detailed information of all the target candidates in the user input DNA sequence. **b** The detailed information of target sites in the reference genome for each target candidate. **c** Visualization of on-and off-target sites with genomic and transcript annotations in JBrowse.

## Discussion

The CRISPR-Cas9 system has been widely applied in genome editing. More recently, the CRISPR-Cpf1 system was identified as a new powerful tool for genome engineering [19]. The CRISPR-Cpf1 system is a class 2 CRISPR-Cas system like CRISPR-Cas9, but it has some distinct characteristics. For example, Cpf1 recognizes a T-rich PAM and cleaves double-strand DNA via a staggered cut,resulting in two sticky ends [19]. Compared with CRISPR-Cas9, the CRISPR-Cpf1 system has some advantages, such as higher target specificity in human cells [23] and better performance in multiplex gene editing [21]. Target efficiency and specificity are the two most important aspects for genome editing. However, target efficiency for the CRISPR-Cpf1 system varies among different gRNAs [41]. gRNA characteristics related to target efficiency have not been well studied. Here, we reanalyzed the published CRISPR-Cpf1 gRNAs data [41] and found that many sequence and structural features of gRNAs (e.g., the position-specific nucleotide composition, position-nonspecific nucleotide composition, GC content, minimum free energy, and melting temperature) are correlated with target efficiency. As a machine learning technology, a SVM model was created based on the published gRNAs data to predict target efficiency for any given gRNAs. We have developed the first web service application, CRISPR-DT, to help users design optimal gRNAs for the CRISPR-Cpf1 system by considering both target efficiency and specificity. The target efficiency score is available for mammals because the SVM model was built based on mammalian data [41]. We have updated our previously published three CRISPR-Cas systems (Cas9, Cas9n, and RFN) by incorporating Doench et al.’s model to predict target efficiency for mammals[54, 56], which are also available in CRISPR-DT. CRISPR-DT will empower researchers in genome editing.

## Methods

### Data retrieval and usage

The published 1251 gRNAs data was downloaded from the journal’s website (https://www.nature.com/articles/nmeth.4104) [41]. The top 10% (125 efficient) and bottom 10% (125 inefficient) gRNAs in activity ranking of the 1251 gRNAs were analyzed to explore features related to target efficiency and were also used to train the SVM model. In addition, the 104 independent gRNAs data for validating the SVM model was downloaded from the journal’s website [41]. The reference genomes and corresponding gff3 annotation files for predicting target specificity were downloaded from Ensembl (https://useast.ensembl.org) and Ensembl Plants (https://plants.ensembl.org) with release 81 for animals and release 27 for plants.

### Computational tools and statistical analysis

The minimum free energy of each 23-nt gRNA sequence was calculated by RNAfold with default parameters [57]. The Melting temperatures of the entire 23-nt gRNA sequence, seed region, trunk region, and promiscuous region were calculated using the Biopython Tm_NN function with thermodynamic values from the DNA_NN2 table [58–60]. Welch’s t-test was used to perform statistical significance analysis (*P*-value < 0.05) by the R package.

### Target efficiency predictive model

The top 10% (125) gRNAs in activity ranking of the 1251 gRNAs were labeled by “efficient”,while the bottom 10% (125) gRNAs were labeled by “inefficient”. We calculated 796 features for the 250 gRNAs, including the position-specific nucleotide composition (23 × 4 + 22 × 4^2^), position-nonspecific nucleotide composition(4 + 4^2^ + 4^3^ + 4^4^ + 1 + 1), GC content (4 + 1), minimum free energy (1), and melting temperature (4). Since the full set of gRNA features was overdetermined, feature selection was preformed carefully using Random Forest before training SVM models. We used ten-fold cross validation to evaluate the SVM model performance. Specifically, the 250 efficient and inefficient gRNAs were randomly divided into ten folds; each fold was used once as the test data and correspondingly, the remaining nine folds were used as the training data. For each time, first, we used Random Forest to select important features (48 features were selected according to the mean decrease impurity) based on the training data; second, optimal parameters of the SVM model were selected following the protocol recommended by LIBSVM [61]; third, we built the SVM model by LIBSVM, using a radial basis function (RBF) as kernel transformation, based on the training data with the important features and optimal parameters; fourth, the SVM model was evaluated by the test data. After repeating the above procedure ten times, we got the average performance of SVM models. The ROCR package of R was used to draw the ROC curve and calculate the AUC value. Last, we built the final SVM model by using all the 250 gRNAs data with the 48 most important features (Additional file 1:Table S3)and optimal parameters. To further validate the final SVM model, we compared our model with previously published model by using 104 independent gRNAs data,in which the top 20% gRNAs in activity ranking were defined as efficient ones and the remaining 80% gRNAs were regarded as inefficient ones [41]. In addition, the 104 gRNAs were divided into three subsets based on different PAM sequences (TTTA, TTTC, and TTTG) to further evaluate the SVM model. In each subset, the top 20% gRNAs in activity ranking were defined as efficient ones and the remaining 80% gRNAs were regarded as inefficient ones [41].

### Bioinformatics pipeline for improving target specificity

The pipeline is shown in Additional file 1: Figure S4. First, users input a DNA sequence. Second, a Perl script was used to search all target candidates from the input DNA sequence based on the on-target PAM sequence users set. Third, each target candidate is mapped to the reference genome by Bowtie2 [62] to find all the possible target sites in the genome according to the setting of maximum number of mismatches and gaps that off targets can tolerate. Fourth, samtools [63] was used to separate alignment results to the forward strand and reverse strand. Fifth, several Perl scripts were used to filter results based on users off-target settings. After converting file format, we use PHP and JavaScript to display the results in DataTables and JBrowse [55]. The same strategy was used in our previous work [56, 64].

## Declarations

### Ethics approval and consent to participate

Not applicable

### Consent for publication

Not applicable

### Availability of data and material

CRISPR-DT is freely accessible via the bioinformatics laboratory website of Miami University [65]. The source code is available from the authors on request.

### Competing interests

The authors declare that they have no competing interests.

### Funding

This project is funded by Office for the Advancement of Research and Scholarship (OARS) and Biology Department, Miami University, Oxford, OH, USA.

### Authors’ contributions

CL managed and coordinated the whole project. HZ and CL participated in CRISPR-DT design. HZ implemented statistical analysis, SVM modeling, and the backend pipeline in Perl. HZ and CL participated in web interfaces implementation. HZ and CL prepared the manuscript. All authors have read and approved the final manuscript.

## Acknowledgements

This project is funded by Office for the Advancement of Research and Scholarship (OARS) and Biology Department, Miami University, Oxford, OH, USA.

## Additional file

**Additional file 1:** Supplementary Figures and Tables. (PDF 572 kb)

